# The 5-HT_1A_ receptor selective agonist NLX-204 displays analgesic activity in the knee osteoarthritis and plantar incisional post-operative pain models in rats

**DOI:** 10.64898/2025.12.02.691847

**Authors:** Ronan Depoortère, Adrian Newman-Tancredi

**Author notes:** Corresponding author: Depoortère R., Neurolixis SAS, 81100 Castres, France.

## Abstract

**Background:** NLX-204 is a highly selective, high efficacy biased agonist at serotonin 5-HT1A receptors, which are well-established modulators of pain processing.

**Objective:** To evaluate the analgesic activity of NLX-204 in two rat models of nociceptive/inflammatory pain: the monoiodoacetate (MIA)-induced knee osteoarthritis (KOA) and the Brennan’s plantar incision model, using morphine as an active comparator, in both male and female rats.

**Methods:** KOA was induced by left intra-articular MIA injection. Analgesic effects of NLX-204 (0.1-3 mg/kg p.o.) or morphine (6 mg/kg s.c.) were assessed on days 5 and 8 by measuring withdrawal threshold (WT) using von Frey filaments and dynamic weight bearing (DWB) on injured and contralateral limbs. In the Brennan’s model, the plantar surface of the left hind limb was incised with elevation and incision of the plantaris muscle. Mechanical allodynia (WT) and guarding behavior (GB) were assessed 24 h post-surgery.

**Results:** In the MIA-KOA model, NLX-204 significantly attenuated DWB imbalance in males at 1-3 mg/kg (acute) and 0.1-3 mg/kg (repeated dosing), and in females at 3 mg/kg (repeated dosing only). NLX-204 demonstrated anti-allodynic activity (VF filaments) from 0.3 mg/kg in males (acute and repeated) and from 1 mg/kg in females. In the Brennan’s model, NLX-204 reduced GB scores at 1 mg/kg (males) and 0.3-1 mg/kg (females), and showed anti-allodynic effects from 0.3 mg/kg in both sexes. Morphine was effective in both models under acute and repeated administration.

**Conclusions:** Oral NLX-204 demonstrates dose-dependent analgesic and anti-allodynic activity in two rat models of nociceptive/inflammatory pain, supporting the therapeutic potential of selective, high-efficacy 5-HT1A receptor agonists as a novel non-opioid strategy for pain management.

## INTRODUCTION

Chronic pain is a widespread pathology that, in addition to causing personal suffering, presents a major public health challenge and substantial economic burden due to direct health care costs and lost productivity. Thus, in the USA alone, the annual cost of pain disorders is estimated at between $560 and $635 billion (2010 equivalent US$), far above the costs of other main pathologies (Gaskin and Richard, 2012). Although there are numerous therapeutic options available to alleviate painful conditions (monoamine reuptake inhibitors, opioids, NaV channel blockers, gabapentin-likes, NSAIDs, etc.), they present various side-effects that can seriously limit their protracted use. For instance, NSAIDs use can lead to gastrointestinal bleeding, renal toxicity and increased cardiovascular risks, while opioids use is associated with a multiplicity of side-effects, such as constipation, respiratory distress, and even death from overdosing. Moreover, opioids possess marked abuse and addiction liability in a substantial proportion of patients. In the so-called “opioid crisis” in the US, opioids caused around 100.000 deaths during the 12-month period ending in April 2021 (https://www.cdc.gov/nchs/pressroom/nchs_press_releases/2021/20211117.htm).

These observations have spurred the search for novel analgesic compounds that would combine an efficacy at least similar to that of NSAIDs, ideally reaching that of opioids, but with a superior side-effects profile. A promising pharmacological target to achieve such a profile is activation of serotonin (5-hydroxytryptamine, 5-HT) 5-HT_1A_ receptors. Indeed, agonists at 5-HT_1A_ receptors have been shown to display analgesic activities in rodents (Colpaert, 2006; Haleem, 2019) and a particular agonist that has been extensively studied in animal pain models is NLX-112 (a.k.a. befiradol or F 13640), which is exceptionally selective for 5-HT_1A_ receptors, possesses high potency in vivo, and displays no affinity for any of the subtypes (μ, δ, and κ) of opioid receptors (Colpaert et al., 2002). In particular, NLX-112 is robustly active in nociceptive, neuropathic and traumatic pain models in rats and mice (Colpaert, 2006) (see (Newman-Tancredi et al., 2022) for review).

Using NLX-112 as a chemical scaffold, we recently synthetized a close chemical congener, NLX-204 (Sniecikowska et al., 2019) which is also highly selective for the 5-HT_1A_ receptor, is highly potent in vivo, and displays no affinity for opioid receptors (Sniecikowska et al., 2019). In addition, NLX-204 possesses favorable pharmacokinetic characteristics (notably high central nervous system penetration) suitable for pharmaceutical targeting of central (spinal cord and brain) 5-HT_1A_ receptors (Sniecikowska et al., 2019). In vivo, NLX-204 possesses marked antidepressant-like activity in the forced swim test, where it potently, efficaciously and dose-dependently reduced immobility (Sniecikowska et al., 2019). Moreover, in the chronic mild stress (CMS) procedure in rats, it opposed CMS-induced decrease of sucrose intake (indicating antidepressant-like activity), attenuated anxiety (elevated plus maze test), and reduced working memory deficits (novel object recognition test) (Newman-Tancredi et al., 2022b; Papp et al., 2023; Papp et al., 2024). However, the activity of NLX-204 in models of pain has not previously been reported, so the present study characterized the analgesic properties of NLX-204 using two rat models of nociceptive/inflammatory pain, the monoiodoacetate (MIA)-induced knee osteoarthritis (KOA) model, and the Brennan’s post-operative (skin incisional) pain model. Of note, anxiety, depression and cognitive deficits are often associated with pain, especially chronic pain, so a drug candidate that combines anxiolytic, antidepressant, pro-cognitive and analgesic activity would be highly desirable. The MIA-KOA model exhibits histopathological changes similar to those observed in human osteoarthritis (de Sousa Valente, 2019), and is also suitable for testing new therapeutic agents for other inflammatory-mediated painful conditions. The Brennan’s model (Brennan, 2011) consists of incising the skin of the plantar aspect of the hindlimb, and elevating and incising the underlying plantaris muscle. This post-operative pain model presents the advantage of being technically straight-forward, triggering a mild localized pain lasting several days, and being responsive to a wide array of clinically used analgesics.

There is a sex-biased difference in both clinical and experimentally induced pain, with female subjects at greater risk for developing several chronic pain disorders and exhibiting greater sensitivity to noxious stimuli in the laboratory, as compared with males (Fillingim et al., 2009; Mogil, 2012). For that reason, we included both males and female rats in the present study; in addition, morphine was included as a positive control in all experiments.

## MATERIALS and METHODS

### Animals

Sprague Dawley male and female rats (Envigo) weighing 200-250 g (MIA-KO) and 180-250 g (Brennan’s) upon arrival were housed in (26 x 47.6 x 20.3 cm^3^) cages (3 of the same sex per cage) containing BedO’Cob^®^ bedding with Nylabones^®^ for enrichment. Rats were acclimated to the facility for a minimum of 7 days prior to initiation of the study, and all animals were examined, handled, and weighed prior to initiation of the study to assure adequate health and suitability. During the study, 12/12 h light/dark cycles were maintained with lights on at 6 am. The room temperature was maintained between 20 and 23 °C with a relative humidity at approximately 50 %. Chow (Lab diet 5001, LabDiet, St. Louis, MO, USA) and water were provided *ad libitum* for the duration of the study. All experimental procedures were performed during the animal’s light cycle phase. Rats were randomly assigned to the various experimental groups and animals from different experimental groups were housed together. Sexes were separated in distinct colony rooms. All housing and testing procedures were implemented in AAALAC-accredited facilities, in accordance with the Principles of Laboratory Animal Care and the approval of the PsychoGenics Inc., Institutional Animal Care and Use Committee.

### The monoiodoacetate (MIA) knee osteoarthritis (KO) model

Rats were first tested for baseline (prior to a knee injection of MIA) dynamic weight bearing (DWB) responses on the left and right hind paws using the Dynamic Weight Bearing Instrument 2.0 (Bioseb). One day after recording of Pre-MIA values, rats received an intraarticular injection of MIA (3 mg/50 μl saline) into the left hindlimb knee joint (Day 0). Weight bearing deficits on the left hind paw were assessed in all rats on Day 4 (Post MIA), and rats were included in the study if they met the following inclusion criterion: the left hind paw weight bearing had to be ≤ 40 % of the total hind paw weight bearing. Rats that met the criterion were then assigned to separate groups based on their Post-MIA score such that group averages were balanced.

### The Brennan’s surgical pain model

#### Surgery

The plantar surface of the left hind-limb was first swabbed generously with Betadine and 70% alcohol. Under isoflurane anesthesia (1-3% isoflurane with 1.5L O_2_), a 1.2 to 1.5 cm incision was made using a #10 blade through the skin and fascia of the plantar aspect of the foot starting 0.5 cm from the proximal edge of the heel and extending towards the toes. Using curved forceps, the plantaris muscle was elevated and incised about 0.8-1.0 cm longitudinally leaving the origin and insertion intact. After hemostasis, the skin was closed with 2 horizontal mattress sutures using 5-0 nylon on an FS-2 needle. Sutures were secured with VetBond^®^ surgical adhesive. Following surgery, the rats recovered in a warmed cage (43°C) and were then single-housed for the duration of the study.

#### Assessment of mechanical allodynia using von Frey filaments

Paw mechanical withdrawal threshold (WT) was assessed three times prior to surgery, at 24 h post-surgery, and at 1, 2, 4, and 6 h after compound administration.

On the day of testing, rats were placed in acrylic white rectangular observation chambers (20.3w x 20.3d x 25.4h cm^3^) with black acrylic lids on an elevated, wire mesh platform and allowed to acclimate for a minimum of 30 min before each session. Paw mechanical WT were assessed as described by Chaplan and colleagues. (Chaplan et al., 1994). Briefly, von Frey filaments (3.61 (0.41 g), 3.84 (0.69 g), 4.08 (1.20 g), 4.31 (2.04 g), 4.56 (3.63 g), 4.74 (5.49 g), 4.93 (8.51 g) and 5.18 (15.14 g), Semmes-Weinstein, Stoelting, Wood Dale IL, USA) were applied to the plantar surface of the hind paw and held for approximately 6-8 s, beginning with the 2.04 g filament. A positive response was noted when the paw was sharply withdrawn. The 50% withdrawal threshold was determined using the Dixon “up-down” method where 4 filament applications following the first change in response were performed (Chaplan et al., 1994):

50% withdrawal threshold (g) = (10[Xf+k*d])/10000)

- “Xf” is defined as the final filament used (in log units).
- The parameter “k” is from the Appendix 1 table in Chaplan et al. (1994) [modified from Dixon et al. (1980)] and will be determined by the pattern of the von Frey responses (e.g. OXOXOX) where ‘O’ indicates no response and ‘X’ denotes a withdrawal;
- The parameter “d” is the difference between stimuli described in Chaplan et al. (1994)

and is a constant: d=0.224.

A maximum of 9 filament applications were used to determine the 50% withdrawal threshold, and based on (Chaplan et al., 1994), animals that did not respond to any filament were assigned a threshold of 15.0 g, while animals that responded to all filaments were assigned a threshold of 0.25 g. WT was measured for both the left and right hind paws, and assessments were made by a blinded investigator.

#### Assessment of hind paw guarding behavior

Rats were acclimated to test chambers for up to 60 min and then scored for guarding behavior. Sixty min following treatment, guarding behavior was scored at 1 h (0 - 60 min), 2 h (61 - 120 min), 4 h (241 - 300 min), and 6 h (360 - 420 min). A guarding score was recorded for each rat every 5 min for 60 min. The scores for each rat were added, and a final score was recorded (max score 39). The following scoring system was used:

0: Rats have the ipsilateral (injured) hind paw flat and are putting weight on it; the hind paw is directly under the body. The plantar surface, toes, and heel of the ipsilateral paw must be on the mesh floor firmly, balancing weight uniformly with the contralateral hind paw.

1: The heel of the ipsilateral hind paw is raised, but the rat is still bearing some weight on the plantar surface and toes. At this stage, the distribution of weight between the hind paws ceases to be equal, and no signs of blanching, or whitening, of the injured hind paw should be a sign of such a case.

2: The ipsilateral hind paw is not bearing weight (may be flat and held out to the side), or the rat is only on its toes. The heel, and possibly the plantar surface and base of the toes are raised up, but not totally away from the mesh floor, with an unbalanced support of weight.

3: The entire injured hind paw is completely off the surface and being held close to the body.

Guarding score was assessed prior to and 24 h post-surgery. A guarding score ≥10 at 24 h post-surgery was required for inclusion in the study. The groups were balanced by body weight and guarding score at 24 h post-surgery prior to pharmacological treatment.

### Pharmacological compounds

NLX-204 [(3-Chloro-4-fluorophenyl)(4-fluoro-4-(((2-(pyridin-2-yloxy)ethyl)-amino)methyl)piperidin-1-yl)methanone, fumarate salt] was provided by Neurolixis, dissolved in water and administered orally (p.o.). Morphine sulfate was obtained commercially (Spectrum), dissolved in saline and administered s.c.. Both compounds were administered at a volume of 1 ml/kg body weight, and doses correspond to the weight of the free base.

### Data analysis

Details of statistical analysis are given in the legend of figures. Two-way ANOVAs with the condition (for example: pre-MIA, post-MIA, 1, 2 and 4 h post-treatment) as the within-subjects factor and the pharmacological treatment (vehicle, dose of NLX-204 or morphine) as the between-subjects factor were followed by Dunnett’s post-hoc tests wherever appropriate and reported by statistical symbols in figures. Statistical analyses were implemented with the Prism^®^ V10.3.1 software.

## RESULTS

For both the MIA-KO and Brennan’s model experiments, there were no significant effects of treatment on body weights throughout the study for both sexes: all animals appeared healthy throughout the study with no signs of motor impairment or other morbidities (data not shown).

### NLX-204, given acutely and repeatedly, attenuates MIA-induced weight bearing deficit in the knee osteoarthritis model

Out of a total of 72 male and 72 female rats assigned to this study, 19 males and 12 females failed to meet the %DWB inclusion criterion.

Under control conditions, i.e., one day before MIA intra left knee injection, male rats presented a % left DWB around 50%, i.e., balanced their weight equally between the two hindlimbs (1^st^ set of bars, upper panel, Fig. 1). Four days after MIA injection, DWB of the left hind paw was drastically reduced to around 20% of total left and right DWB (2^nd^ set of bars), indicating that rats favored the uninjured (right) hindlimb for weight bearing. One day after MIA injection, the DWB score of rats injected with saline (30 min pre-test) remained low, at 25% (black bar of third set of bars). By contrast, rats injected acutely with NLX-204 displayed a dose-dependent increase in the %DWB of the left hindlimb, with significant effects versus saline treatment from 1 mg/kg p.o., peaking at 36%, a value quite close to that of morphine, used as an active comparator (40% at 6 mg/kg s.c.). The beneficial activities of NLX-204 and morphine were preserved following repeated treatment (b.i.d. over 3 days) (last cluster of bars).

**Figure 1:**
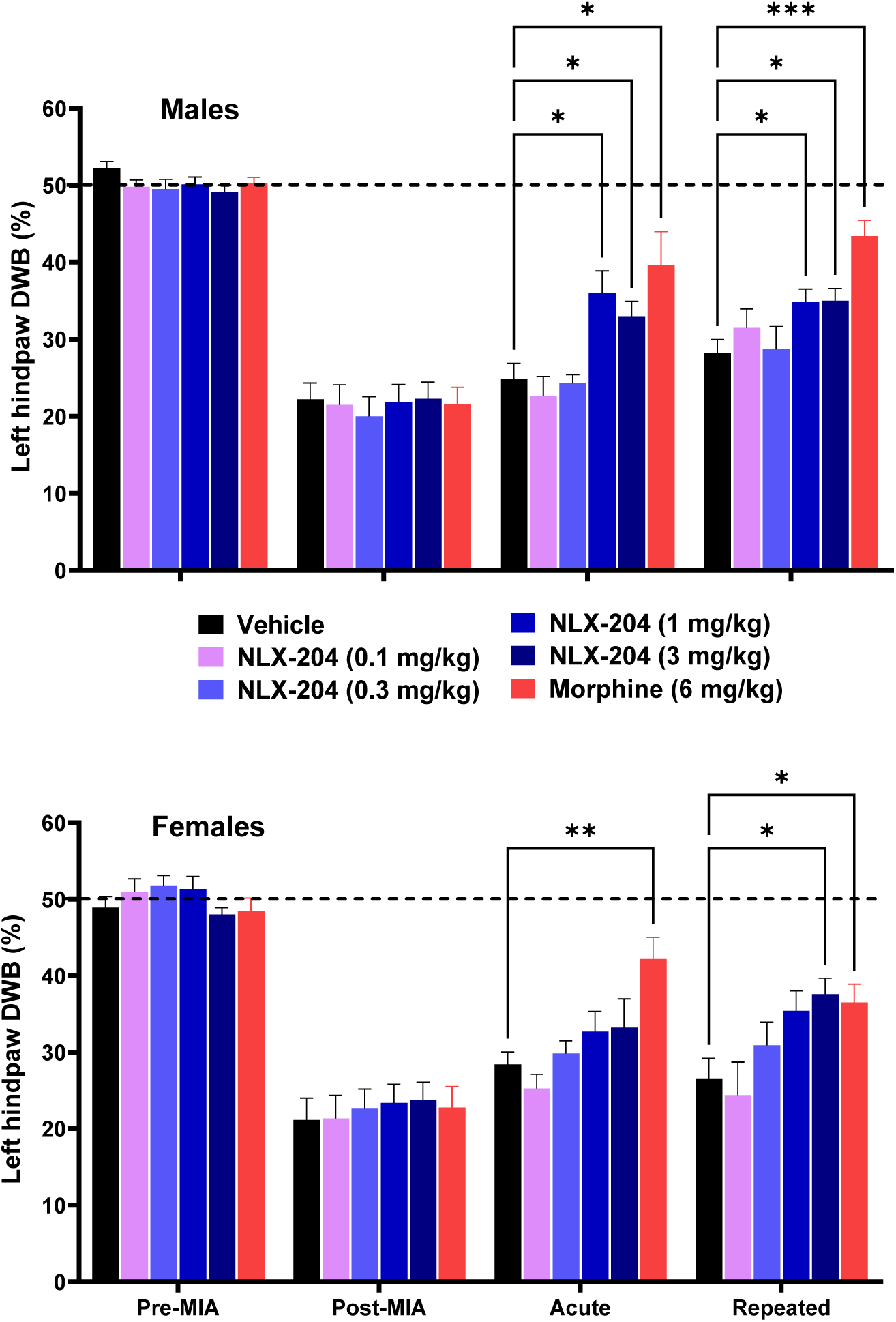
Left hind paw dynamic weight bearing (DWB) in male (top panel) and female (bottom panel) rats prior to and following MIA injection, and effects of vehicle, NLX-204 or morphine, at 1 h post-dosing on Day 5 following a single acute administration, and at 1 h post-dosing on Day 8 following repeated administration. Bars are mean with SEM. N=10 rats per group. * p<0.05, ** p<0.01, *** p<0.001, Dunnett’s post-hoc test following significant two-way ANOVAs. For male rats: F (5, 54) = 6.50; F (2.719, 150.2) = 216.4 and F (15, 162) = 3.72, all P’s<0.0001; for female rats : F (5, 54) = 3.80; F (2.531, 136.7) = 144.2 and F (15, 162) = 2.31, all P’s<0.01, for the treatment, time and treatment x time factors, respectively.

Results with female rats followed a pattern on the whole similar to that of male rats, except that NLX-204 appeared to be less potent, with a MED dose of 3 mg/kg observed following repeated treatment (lower panel, Fig. 1). Of note, morphine also appeared to be less potent in females than in males.

### NLX-204, given acutely, attenuates MIA-induced mechanical allodynia in the knee osteoarthritis model

Before MIA intra-knee injection, the vast majority of male rats withstood application of the 15.0 g VF filament without withdrawal of the left (i.e. injured) hindlimb (1^st^ set of bars, upper panel, Fig. 2). Five days following MIA injection, rats withdrew their hindlimb following application of VF filaments with an average of 1.2 g (2^nd^ set of bars), i.e., demonstrated marked mechanical allodynia. One h following acute administration, NLX-204 significantly increased the force that could be applied before withdrawal, i.e., attenuated mechanical allodynia, from 0.3 mg/kg onwards. The attenuating effect of NLX-204 still appeared to be present at 2 h post-treatment, although not significant, likely due to a fairly high dispersion of values.

**Figure 2:**
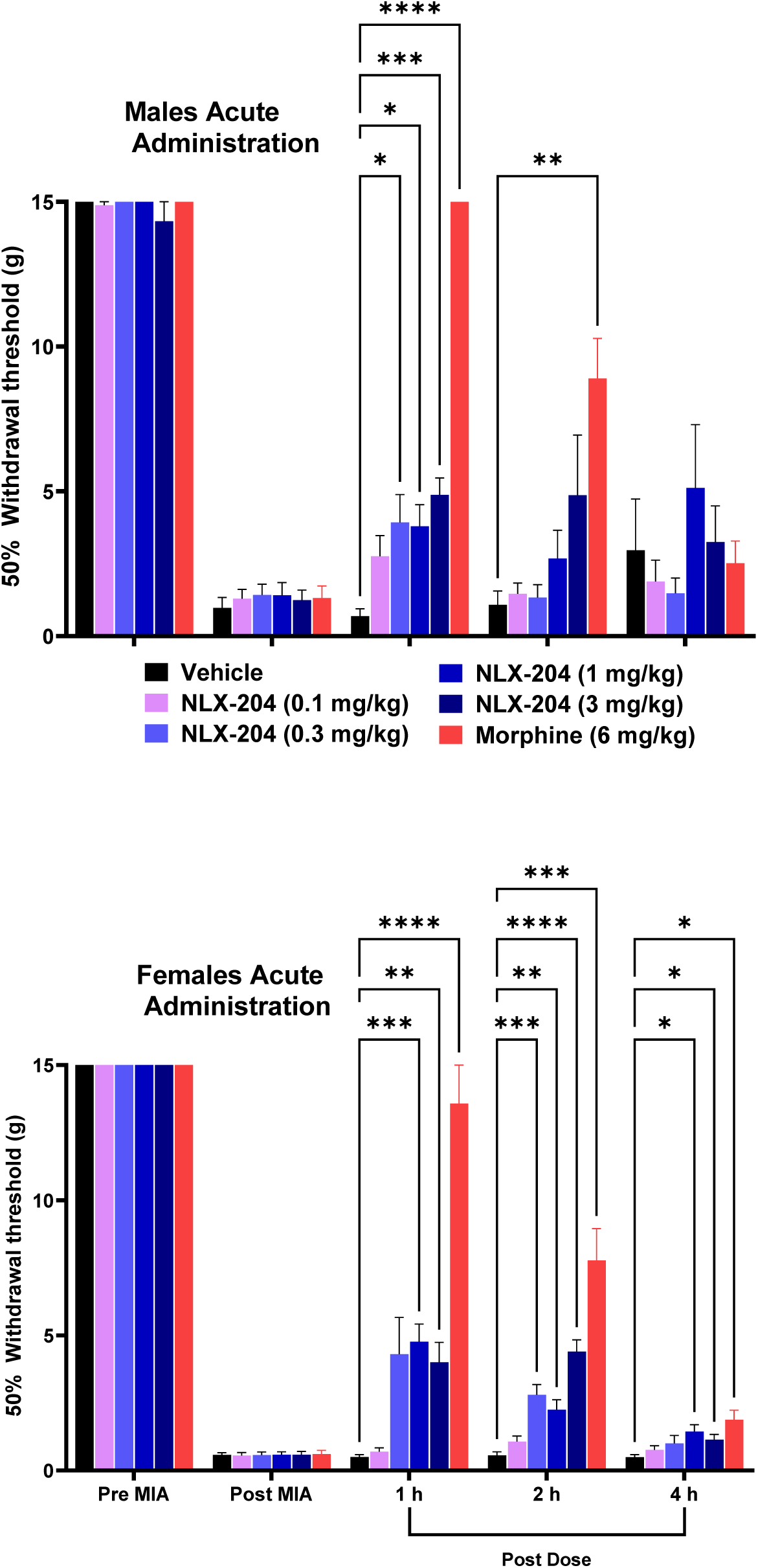
Left hind paw tactile sensitivity (VF filaments) in male (top panel) and female (bottom panel) rats prior to and following MIA injection, and effects of vehicle, NLX-204 or morphine at 1, 2, and 4 h post-dosing on Day 5 following a single acute administration. Bars are mean with SEM. N=8 to 10 rats per group; * p<0.05, ** p<0.01, *** p<0.001 **** p<0.0001, Dunnett’s post-hoc test following significant ANOVA. For male rats: F (5, 47) = 8.92; F (2.300, 108.1) = 308.9 and F (20, 188) = 10.25, all P’s<0.0001; for female rats : F (5, 54) = 53.73; F (1.796, 96.96) = 852.0 and F (20, 216) = 18.24, all P’s<0.0001, for the treatment, time and treatment x time factors, respectively.

In female rats, surgery-induced mechanical allodynia was a bit more pronounced than that in males (on average 0.5 g for withdrawal); the attenuating effects of NLX-204 lasted longer than in males, with significant effects still observed at 4 h at 1 and 3 mg/kg.

Morphine, in both sexes, was more efficacious than NLX-204, with a time-dependent decrease in its pharmacological activity.

After surgery, WT for the contralateral (uninjured) hind paw were recorded as a maximum of 15.0 g throughout the study for all groups, males or females (data not shown).

### NLX-204, given repeatedly, attenuates MIA-induced mechanical allodynia in the knee osteoarthritis model

On day 8, following twice-daily treatments starting on day 5, NLX-204 significantly increased WT at 3 mg/kg and 1 h post-dosing in both males (upper panel, Fig. 3) and females (lower panel). It displayed a trend to increase WT at lower doses for all three times post-dosing. Morphine was active at all three times (males) and at 1 and 2 h (females) post-dosing

**Figure 3:**
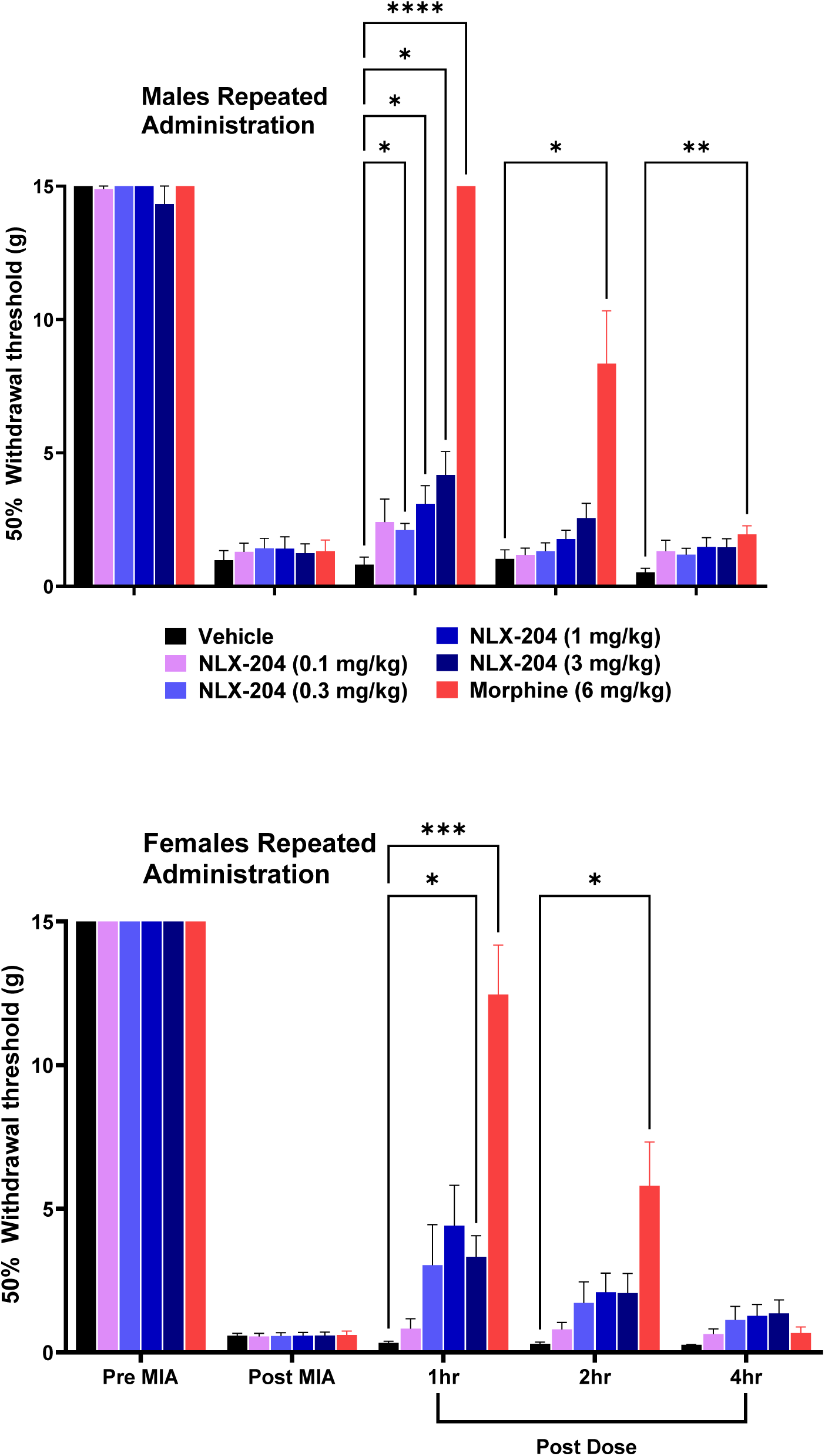
Same as in Figure 2, except that the effects of vehicle, NLX-204 or morphine were assessed at 1, 2, and 4 h post-dosing on Day 8 following 3 days of repeated administration. Bars are mean with SEM. N=8 to 10 rats per group; * p<0.05, ** p<0.01, *** p<0.001 **** p<0.0001, Dunnett’s post-hoc test following significant ANOVA: For male rats: F (5, 47) = 43.31; F (2.367, 111.2) = 675.1 and F (20, 188) = 18.47, all P’s<0.0001; for female rats : F (5, 54) = 12.23; F (1.695, 91.51) = 636.5 and F (20, 216) = 11.06, all P’s<0.0001, for the treatment, time and treatment x time factors, respectively.

### NLX-204, given acutely, diminishes mechanical allodynia in the Brennan’s surgical pain model

Before surgery, the WT for the ipsilateral (surgery) side remained maximal for all male rats (15.0 g VF filament: 1^st^ set of bars, upper panel Fig. 4). Twenty four h following surgery, the value dropped to the minimum, i.e. they all withdrew with application of the finest (0.4 g) filament (2^nd^ set of bars), indicating marked mechanical allodynia. NLX-204 dose-dependently attenuated this drop of WT (i.e., mechanical allodynia), with significant effects observed at 0.3 and 1 mg/kg p.o. at 1 and 2 h post-administration. Indeed, at 1 h post-administration, NLX-204 1 mg/kg almost abolished the surgically-induced drop in WT. Morphine (6 mg/kg s.c.) was on the whole as efficacious as 1 mg/kg NLX-204 at all three time periods.

**Figure 4:**
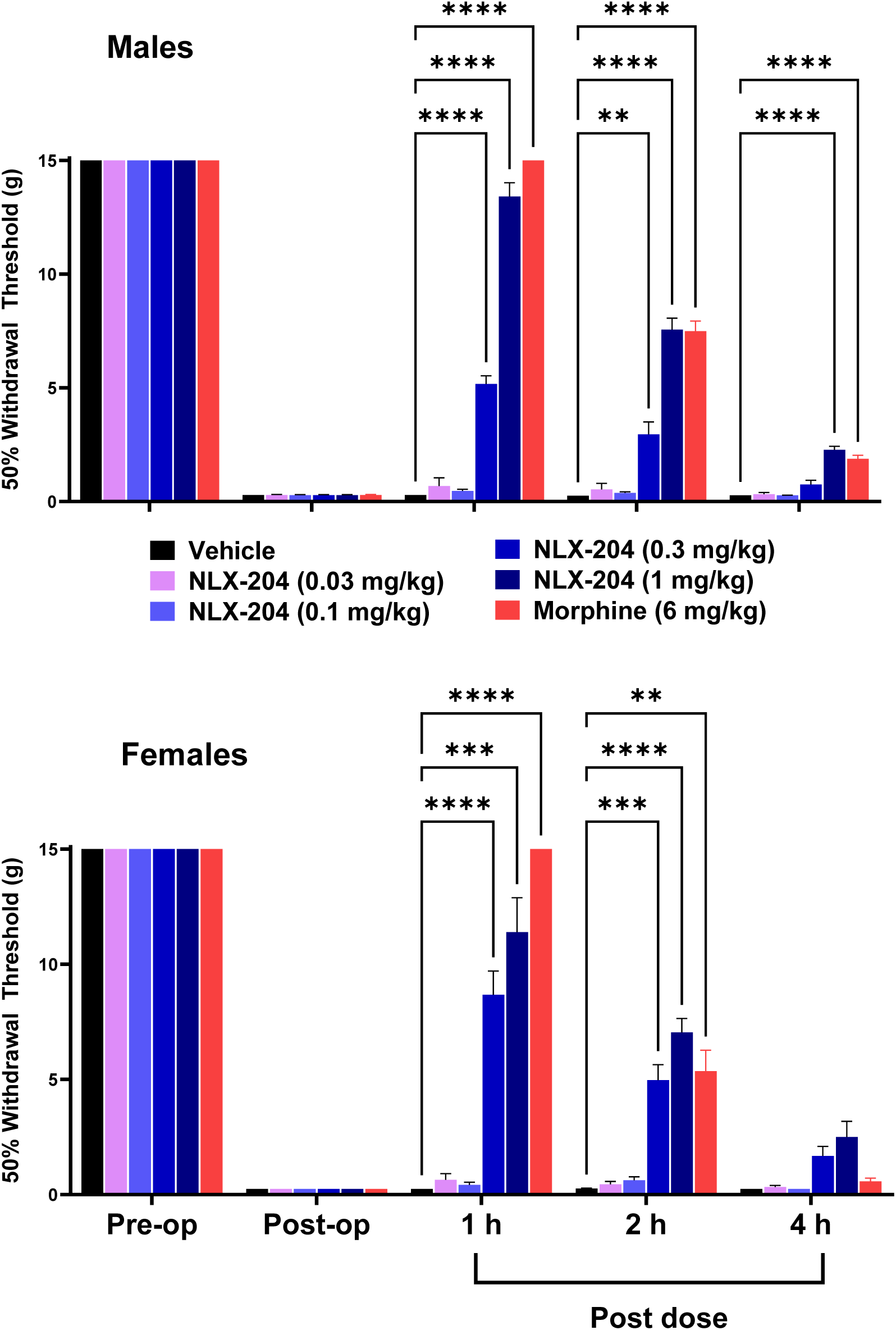
Left hind paw tactile sensitivity (VF filaments) in male (top panel) and female (bottom panel) rats prior to and following plantar incision, and effects of vehicle, NLX-204 or morphine at 1, 2, and 4 h post-dosing 24 h post-surgery following a single acute administration single administration. Bars are mean with SEM. N=10 rats per group; * p<0.05, ** p<0.01, *** p<0.001, **** p<0.0001, Dunnett’s post-hoc test following significant ANOVA. F (5, 54) = 334.1; F (2.098, 113.3) = 4651 and F (20, 216) = 184.5, all P’s<0.0001; for female rats: F (5, 54) = 88.9; F (2.111, 114.0) = 1187 and F (20, 216) = 43.9, all P’s<0.0001, for the treatment, time and treatment x time factors, respectively.

For the cohort of female rats, the results were quite similar, except for the fact that NLX-204 appeared to be a bit more efficacious than for male rats at 0.3 mg/kg (lower panel, Fig. 4).

After surgery, WT for the contralateral (uninjured) hind paw were recorded as a maximum of 15.0 g throughout the study for all groups, males or females (data not shown).

### NLX-204, given acutely, attenuates guarding behavior in the Brennan’s post-operative pain model

Guarding behaviors were not observed in any male rats prior to surgery (score of 0, not visible, 1^st^ set of bars of upper panel, Fig. 5) but were observed in all groups following surgery (maximal score of 39, 2^nd^ set of bars). NLX-204 attenuated the severity of guarding behaviour, with significant effects at 1 mg/kg at 1 and 2 h post-administration. Unlike what was observed for WT above, morphine was more efficacious than NLX-204, particularly at 1 and 2 h post-treatment.

**Figure 5:**
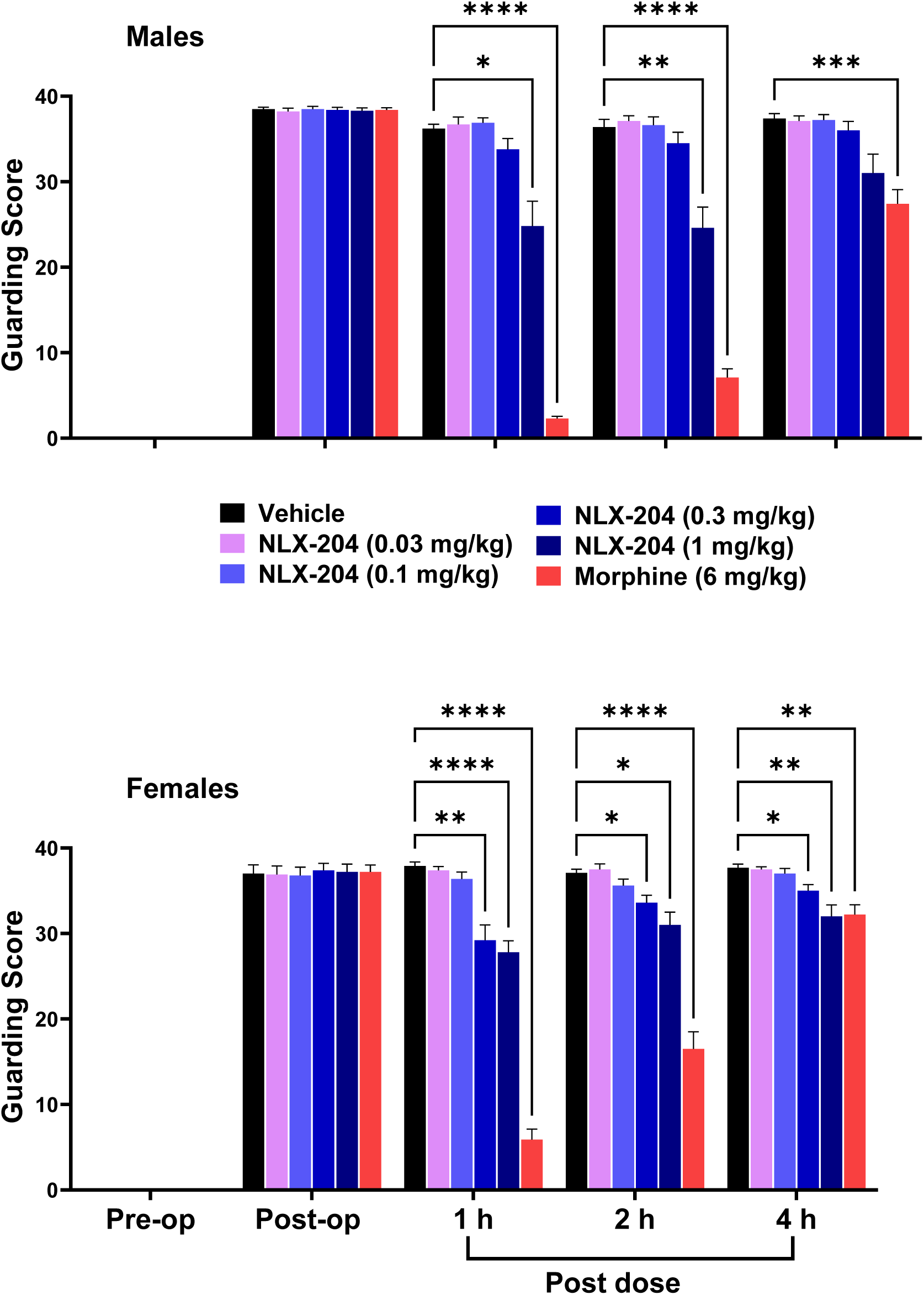
Left hind paw guarding score in male (top panel) and female (bottom panel) rats prior to and following plantar incision, and effects of vehicle, NLX-204 or morphine at 1, 2, and 4 h post-dosing 24 h post-surgery following a single acute administration single administration. Bars are mean with SEM. N=10 rats per group; * p<0.05, ** p<0.01, *** p<0.001 **** p<0.0001, Dunnett’s post-hoc test following significant ANOVA. F (5, 54) = 67.0; F (2.661, 143.7) = 1904 and F (20, 216) = 58.6, all P’s<0.0001; for female rats : F (5, 54) = 71.8; F (3.288, 177.6) = 1948 and F (20, 216) = 42.1, all P’s<0.0001, for the treatment, time and treatment x time factors, respectively.

In female rats, a set of data similar to that of male rats was recorded (lower panel, Figure 5), and in accord with what was observed for WT, NLX-204 appeared to be a bit more efficacious in this sex, with significant effects at 0.3 mg/kg from 1 to 4 h.

## DISCUSSION

The main findings of the present study are as follows: 1) NLX-204 displayed dose-dependent analgesic activity in the MIA-induced KOA and Brennan’s post-surgical pain models; 2) the analgesic activity of NLX-204 was maintained following repeated dosing in the former model 3) both male and female rats responded (at various doses) to the analgesic activity of NLX-204 in both pain models.

### NLX-204 is analgesic against osteoarthritis-induced and surgically-induced pain

Osteoarthritis, a pathology with a strong inflammatory component (Berenbaum, 2013) is a major cause of chronic pain, with the knees as the most common sites, and is considered to affect close to 600 million people worldwide, with a prevalence anticipated to rise by 60-100% by 2050 (Collaborators, 2023) and which therefore constitutes a substantial medical and societal challenge. Here, In the MIA-KO model, NLX-204 produced dose-dependent analgesic effects following both acute and repeated administration. Thus, when NLX-204 was administered repeatedly (twice daily for 3 days, starting on day 5), its analgesic activity was preserved when tested on day 8, both for DWB and for the VF assessment of hyperalgesia, suggesting and absence of tachyphylaxis to the analgesic activity of the compound. Of note, the efficacy of 3 mg/kg NLX-204 was equivalent to that of morphine in female subjects. This is of prime importance, as osteoarthritis is a chronic condition necessitating protracted administration of pain killers. Studies using longer treatment periods in the MIA-KO and other pain models are warranted to further validate the present findings.

As concerns post-surgical pain, it is not properly dealt with in greater than 80% of surgery patients in the US, although rates vary depending on such factors as type of surgery performed, and the analgesic/anesthetic intervention used. Inadequate control of post-surgical pain leads to numerous problems, such as delayed recovery, limited physical function, impaired quality of life, surge in cost of care and even increased morbidity (Gan, 2017). In addition, inadequate control of early postoperative pain can lead to chronicity of pain for months after surgery in a substantial proportion of patients (Fregoso et al., 2019). There is hence great need for non-opioid pharmacotherapy to control pain following surgery.

In the present study, both guarding behavior and mechanical allodynia were dose-dependently diminished by NLX-204 at 24 h following surgery. In the case of mechanical allodynia the effects of NLX-204 were more marked, with the dose of 1 mg/kg being nearly as efficacious as 6 mg/kg morphine in male rats. Similarly, NLX-112, a.k.a. F 13640 or befiradol (as a reminder, a close chemical congener of NLX-204), was found to be potently analgesic in a variant of the Brennan’s model of surgical pain (Kiss et al., 2005).

Of note, data from human studies support the translational validity of preclinical incisional pain models. Hence, an incision practiced in the forearm of human volunteers triggered mechanical hyperalgesia (assessed with VF hairs) (Kawamata et al., 2002a; Kawamata et al., 2002b) that was similar in magnitude and duration to the mechanical hyperalgesia in rats after plantar incision. Further, local activation of 5-HT_1A_ receptors with a topical cream containing 8-OH-DPAT produced smaller wound area, scar size, and improved neovascularization, all of which contributed to improve healing outcomes in an excisional punch biopsy study in mice (Sadiq et al., 2018). Thus, the use of a 5-HT_1A_ receptor agonist could promote wound healing, in addition to analgesic activity.

### The analgesic activity of NLX-204 is congruent with that of other 5-HT_1A_ receptor agonists

The serotonergic system has long been implicated in the control of pain processing (Sommer, 2004; Wei et al., 2012): for example, antidepressants, whose primary mode of action is to augment the serotonergic tone, have notable analgesic activity (Bajwa et al., 2009). At the clinical level, tricyclic antidepressants are recognized as having analgesic efficacy independently of their effects on the underlying depression (Janakiraman et al., 2016). Regarding the receptor subtypes most likely to be involved in the analgesic activity of serotonergic compounds, the 5-HT_1A_, 5-HT_3_ and 5-HT_7_ subtypes are pre-eminent targets. The 5-HT_1A_ receptor, in particular, has been the object of numerous preclinical pain studies, using older agonists such as 8-OH-DPAT or buspirone (Bardin, 2011; Haleem et al., 2018) or the more recent and highly selective agonist, NLX-112 (Colpaert, 2006).

### The analgesic activity of NLX-204 in both pain models is specific, and not due to sedative and/or motor interference

Some behavioral readouts for the two pain models rely on measures of reflexive behaviors such as paw WT (von Frey hairs) and guarding behavior, and could therefore be sensitive to sedative and/or motor incapacitating effects. However, this is unlikely to be the case for NLX-204: at all three doses tested, it elicited hardly any effects on two relevant motor items of the Irwin test, locomotor activity and sedation/excitation (supplementary Table 1). In addition, NLX-204 triggered ataxia in some rats at 1 h but not 2 h, although NLX-204 was analgesic at both time points.

### The analgesic activity of NLX-204 is observed in both male and female rats

The study used rats of both sexes, since as alluded to in the Introduction, female subjects are at greater risk for developing chronic pain (Fillingim et al., 2009; Mogil, 2012; Ruau et al., 2012). However, females have been mostly excluded from preclinical experiments on pain (Mogil and Chanda, 2005; Mogil, 2012), based on the commonly held assumption that the estrous cycle and its associated hormonal swings complicate pain preclinical studies and may confound their results. In the present study, NLX-204 exerted analgesic effects in both males and female rats, although significance was achieved in males at lower doses than females in the MIA model, whereas the opposite was true for the Brenan’s model. Overall, it may be concluded that both males and females are sensitive to selective activation of 5-HT_1A_ receptors. Although there does not seem to be comparative data between sexes for the antinociceptive activities of 5-HT_1A_ receptor agonists, it is generally the case that, for unclear reasons, morphine is more potent in males than females in preclinical pain models (Baamonde et al., 1989; Barrett et al., 2001). To summarize this comparison between sexes, the present data suggest that NLX-204 might be efficaciously analgesic in female subjects, which constitutes a definite advantage, considering that chronic pain disproportionately affects more women than men (Mogil, 2012).

### Possible involvement of descending brain stem to spinal cord serotonin pathways in the analgesic activity of NLX-204

In the rat, 5-HT_1A_ receptors are expressed in various brain regions, including the frontal cortex, brain stem (raphe nuclei) and spinal cord (Pompeiano et al., 1992). Whereas the frontal cortex is involved in the antidepressant-like effects of 5-HT_1A_ receptor agonists (Newman-Tancredi et al., 2018), the spinal cord is implicated in their analgesic effects (Kim et al., 2015; Nadeson and Goodchild, 2002). There is indeed a direct serotonergic projection from the midbrain Raphe nuclei to the spinal cord (Liu et al., 2002). Hence, intrathecal (i.t.) administration of the prototypical 5-HT_1A_ receptor agonist 8-OH-DPAT in the L5-L6 region of the spinal cord induced analgesic effects in the rat formalin model of tonic nociceptive pain (Bardin and Colpaert, 2004). Using the same experimental protocol, analgesic activity has also been observed using i.t. NLX-112 (Newman-Tancredi et al., 2018). Of note, the effects of NLX-112 were reversed by co-administration of the 5-HT_1A_ receptor antagonist, WAY-100,635. Furthermore, the analgesic effects of systemically administered NLX-112 were also reversed by i.t. administration of WAY-100,635, most notably on paw licking. Taken together, these elements indicate that the spinal cord is most likely a primary locus for the analgesic activity of NLX-204, although formal experiments to verify this hypothesis are warranted.

### Perspectives and Conclusions

The 5-HT_1A_ receptor agonistic activity of NLX-204 potentially offers additional advantages for its use as an analgesic. Indeed, in addition to displaying analgesic activity, NLX-204 possesses other interesting features: recent studies in rodents have shown that NLX-204 also displays marked antidepressant, anxiolytic and cognitive enhancing activities (Papp et al., 2023; Papp et al., 2024; Sniecikowska et al., 2019). Considering that a substantial percentage of patients suffering from chronic pain conditions show co-morbid depression, anxiety and cognitive deficits (Butler, 2017; de Heer et al., 2014; Linton and Bergbom, 2011), the possibility of having a pharmacological approach that can target both pain and the above-mentioned co-morbid pathologies is all the more attractive. These activities were anticipated to be displayed by NLX-204, considering that 5-HT_1A_ receptor agonists are well documented to present these properties (see: (Haleem, 2019; Lacivita et al., 2012) for review).

5-HT_1A_ receptor agonists, in addition to possessing analgesic, antidepressant, anxiolytic and cognitive enhancing activities (see above), present several additional characteristics that could be of interest for an analgesic. First, NLX-112 had fentanyl dose-sparing activity in a rat chronic pain model of Freund’s adjuvant-induced polyarthritis (Colpaert et al., 2002; Depoortere et al., 2024). In extenso, NLX-112 administered chronically via osmotic sub-cutaneous minipumps (0.63 mg/day) decreased fentanyl oral self-administration by 47 %. In contrast, the prototypical 5-HT_1A_ receptor agonist 8-OH-DPAT, which is less efficacious than NLX-204 for activation of 5-HT_1A_ receptors in vitro, (Newman-Tancredi et al., 2017) was ineffective in reducing fentanyl oral self-administration.

Second, opioids such as fentanyl can produce marked sedation and bradypnea, the latter possibly becoming lethal if too severe; NLX-112 (0.2 mg/kg) reduced fentanyl-induced sedation and respiratory depression by about half (Ren et al., 2015). Similarly, the 5-HT_1A_ receptor partial agonists, tandospirone or 8-OH-DPAT, prevented fentanyl or sufentanyl-induced respiratory depression (Dandrea and Cotten, 2021; Fan et al., 2022; Song et al., 2025), an effect inhibited by WAY-100,635 (Song et al., 2025), confirming the implication of 5-HT_1A_ receptors against the bradypneic effects of opioids.

Third, contrary to opioids, 5-HT_1A_ receptor agonists such as buspirone, gepirone or tandospirone are considered to be devoid of abuse potential, based on preclinical data and clinical observations (Balster, 1990; Sannerud et al., 1993). Likewise, NLX-112 is devoid of abuse liability, based on rat intracranial self-stimulation (ICSS) and monkey cocaine discrimination experiments (Depoortere et al., 2022).

It is thus plausible that NLX-204 could, similarly to NLX-112 or other 5-HT_1A_ receptor agonists, not only display opioid dose-sparing effects, but also prevent opioid-induced bradypnea, as well as lack abuse liability of its own. The first property would be highly desirable in the case where abrupt cessation of opioids would be unacceptable, and where a tapering down of opioids with gradual introduction of NLX-204 would be more appropriate.

In conclusion, the present data support the analgesic potential of NLX-204 against osteoarthritis- and surgery-induced pain, two pain conditions with a marked inflammatory component and that would benefit from novel pharmacotherapeutic options. These data indicate that NLX-204 might possess analgesic activity against a wide spectrum of pain modalities. In addition, based on the opioid-sparing effects of 5-HT_1A_ receptor agonists, and in particular of the chemical analog NLX-112, and on their lack of abuse potential, it would be desirable to investigate if this is also the case with NLX-204.

## ACKNOWLEDGEMENTS

This work was supported by the National Institute of Neurological Disorders and Stroke/Preclinical Screening Platform for Pain (NINDS/PSPP) contract n° 75N95019D00026.

