## Supplementary material for "The 5-HT_1A_ receptor selective agonist NLX-204 displays analgesic activity in the knee osteoarthritis and plantar incisional post-operative pain models in rats": Supp. Table 1

Extract from an Irwin test done on 16 rats (8 males and 8 females, not included in the pain studies) for each treatment condition.

| ***Treatment*** | ***Vehicle*** | | | | | ***0.3 mg/kg*** | | | |  | ***1 mg/kg*** | | | | | ***3 mg/kg*** | | | | | ***10 mg/kg*** | | | | |
| --- | --- | --- | --- | --- | --- | --- | --- | --- | --- | --- | --- | --- | --- | --- | --- | --- | --- | --- | --- | --- | --- | --- | --- | --- | --- |
| ***Time post-treatment (h)*** | ***0*** | ***1*** | ***2*** | ***4*** | ***24*** | ***0*** | ***1*** | ***2*** | ***4*** | ***24*** | ***0*** | ***1*** | ***2*** | ***4*** | ***24*** | ***0*** | ***1*** | ***2*** | ***4*** | ***24*** | ***0*** | ***1*** | ***2*** | ***4*** | ***24*** |
| **Locomotor Activity** |  |  |  |  |  |  |  |  |  |  |  |  |  |  |  |  |  |  |  |  |  |  |  |  |  |
| Normal | 16 | 15 | 14 | 16 | 16 | 16 | 16 | 16 | 15 | 14 | 16 | 16 | 16 | 15 | 16 | 16 | 16 | 16 | 16 | 12 | 16 | 14 | 16 | 16 | 14 |
| Increased | 0 | 0 | 0 | 0 | 0 | 0 | 0 | 0 | 0 | 0 | 0 | 0 | 0 | 0 | 0 | 0 | 0 | 0 | 0 | 3 | 0 | 0 | 0 | 0 | 2 |
| Decreased | 0 | 1 | 2 | 0 | 0 | 0 | 0 | 0 | 1 | 2 | 0 | 0 | 0 | 1 | 0 | 0 | 0 | 0 | 0 | 1 | 0 | 2 | 0 | 0 | 0 |
| **Sedation/Excitation** |  |  |  |  |  |  |  |  |  |  |  |  |  |  |  |  |  |  |  |  |  |  |  |  |  |
| Normal | 16 | 15 | 15 | 16 | 16 | 16 | 16 | 16 | 15 | 15 | 16 | 16 | 16 | 16 | 16 | 16 | 15 | 16 | 16 | 14 | 16 | 16 | 16 | 16 | 11 |
| Increased | 0 | 0 | 0 | 0 | 0 | 0 | 0 | 0 | 0 | 0 | 0 | 0 | 0 | 0 | 0 | 0 | 0 | 0 | 0 | 2 | 0 | 0 | 0 | 0 | 5 |
| Decreased | 0 | 1 | 1 | 0 | 0 | 0 | 0 | 0 | 1 | 1 | 0 | 0 | 0 | 0 | 0 | 0 | 1 | 0 | 0 | 0 | 0 | 0 | 0 | 0 | 0 |
| **Ataxia** |  |  |  |  |  |  |  |  |  |  |  |  |  |  |  |  |  |  |  |  |  |  |  |  |  |
| Normal | 16 | 16 | 16 | 16 | 16 | 16 | 16 | 16 | 15 | 16 | 16 | 15 | 16 | 16 | 16 | 16 | 11 | 15 | 16 | 16 | 16 | 9 | 15 | 16 | 16 |
| Increased | 0 | 0 | 0 | 0 | 0 | 0 | 0 | 0 | 1 | 0 | 0 | 0 | 0 | 0 | 0 | 0 | 5 | 0 | 0 | 0 | 0 | 6 | 1 | 0 | 0 |
| Decreased | 0 | 0 | 0 | 0 | 0 | 0 | 0 | 0 | 0 | 0 | 0 | 0 | 0 | 0 | 0 | 0 | 0 | 1 | 0 | 0 | 0 | 1 | 0 | 0 | 0 |

Number of observations for normal, increased or decreased activity for vehicle or NLX-112 in three motor items of the Irwin test.
